# Genome-wide Identification of Disease Resistance Genes (R Genes) in Wheat

**DOI:** 10.1101/2020.07.18.210286

**Authors:** Ethan J. Andersen, Lauren E. Lindsey, Madhav P. Nepal

## Abstract

Proteins encoded by plant resistance genes (R genes) detect pathogenic effectors and initiate immune responses. Although R genes in many plant genomes are identified, they are yet to be identified in wheat. The major objectives of this project were to conduct genome-wide identification of the NB-ARC-encoding R genes in wheat (*Triticum aestivum* L.) and assess their genomic architecture and potential functional divergence. Wheat protein sequences were obtained from the Ensembl Genomes database, and genes were identified using interpro program. Chromosomal locations of the R genes were determined and syntenic analyses were performed. Altogether, 2151 wheat NB-ARC-encoding genes were identified, among which 1298 genes formed 547 gene clusters. Many of these gene clusters included highly similar genes likely formed by tandem duplications. Among the NB-ARC-encoding genes, 1552 (∼72%) encode Leucine-Rich Repeats (LRRs), 802 are Coiled-Coil (CC) domain-encoding CC-NBS-LRR (CNL) genes and three are Resistance to Powdery mildew 8 (RPW8) domain-encoding RPW8-NBS-LRR (RNL) genes. Surprisingly, five of the NB-ARC-encoding genes encoded a Toll/Interleukin-1 Receptor (TIR), with no LRR, known as TN genes. CNL clades formed similar nesting patterns with a large expansion of CNL-C group like previously reported findings in wheat relatives. Comparisons of the wheat genome with barley (*Hordeum vulgare* L.) and Tausch’s goatgrass (*Aegilops tauschii* Coss.), showed similar locations for homologous NB-ARC-encoding genes. These results showed that R genes in wheat have diversified through duplication to encode receptors that recognize rapidly evolving pathogenic effectors. Identified R genes in this study have implications in plant breeding, as a source of resistance for crop improvement.

## INTRODUCTION

Wheat (*Triticum aestivum* L.) provides approximately 20% of the human population’s caloric intake (FAOSTAT, 2013) and is afflicted by over 100 different diseases caused by hundreds of pathogen and pest species (Murray, 2015). Diseases that substantially reduce yield, such as biotrophic rusts and necrotrophic leaf spotting diseases, impact global markets and food supply. Historically devastating pathogens continue to produce new strains, which overcome past sources of resistance (i.e. Ug99) (Saari and Prescott, 1985; Pretorius *et al*., 2000; Huerta-Espino *et al*., 2011; Singh *et al*., 2011). Leaf spotting diseases similarly cause a high percentage of yield loss under favorable conditions and appear as several forms: *Septoria tritici* blotch, spot blotch, *Stagonospora nodorum* blotch, and tan spot (Singh *et al*., 2016). Additional diseases like Fusarium head blight, wheat streak mosaic virus, and powdery mildew also plague wheat fields with symptoms that are difficult and costly to manage. Disease resistant wheat cultivars, such as those bred by Edgar McFadden and Norman Borlaug (McFadden, 1930; Borlaug, 1983), have been a significant contribution to agriculture and the understanding of phytopathology. With the help of advanced genetic and genomic technologies, recent efforts identified resistance genes (R genes) that confer resistance to strains of various pathogens, such as the recently discovered Ug99 R-genes *Sr33* and *Sr35* (Periyannan *et al*., 2013; Saintenac *et al*., 2013). The hexaploid bread wheat genome (AABBDD), a draft of which has recently become available ^10,11^, formed through the hybridization of three separate species: *Triticum urartu* (A), an unknown relative of *Aegilops speltoides* (B), and *Aegilops tauschii* (D)(Jia *et al*., 2013; Ling *et al*., 2013; Marcussen *et al*., 2014). This unique polyploidy may have resulted in novel regulatory mechanisms that were necessary due to the presence of multiple progenitor resistance signaling pathways. The large and redundant nature of wheat’s hexaploid genome makes it a good candidate for studying R gene evolution with respect to the recent polyploidization events. Detailed understanding of wheat immune system components provides a framework for further attempts at the development of durable resistance in cereals necessary for reduced yield loss from biotic stress. R genes are generally receptors encoding a Nucleotide-Binding Site and a Leucine-Rich Repeat, giving them the alternative names NBS-LRR or NLR. The LRR is thought to interact with pathogen effectors, allowing the protein to initiate signaling mechanisms (i.e. kinases and transcription factors) that initiate defense responses. The primary domain found in the NBS is the Nucleotide-Binding site found in Apoptotic protease activating factor 1, R genes, and *Caenorhabditis elegans* death-4 protein (NB-ARC). The NB-ARC is associated with ATP/ADP binding, since the molecule uses this upon activation. The main objectives of this research was to conduct genome-wide identification of wheat NB-ARC encoding R genes and assess their genomic architecture and potential functional divergence with respect to their homologs in the genomes of wheat relatives.

## MATERIALS AND METHODS

Wheat chromosome, gene, and protein sequences were downloaded from the Ensembl Genomes database through the Biomart tool (Kersey *et al*., 2014). InterProScan (Jones *et al*., 2014) was used to identify wheat protein sequences with NB-ARC domains (PF00931). Locations for genes encoding NB-ARCs were analyzed for clusters as described in Jupe et al. (2012) using the criteria that genes be within 200,000 bases of each other with fewer than eight addition genes between them (Jupe *et al*., 2012). NB-ARC domain motifs were also assessed using MEME software(Bailey *et al*., 2009), identifying those with P-loop, Kinase-2, and GLPL motifs. Clustered genes with the aforementioned motifs were aligned and manually curated using ClustalW2 integrated within the program Geneious (Larkin *et al*., 2007; Kearse *et al*., 2012). The program MEGA was used for phylogenetic analysis (Kumar *et al*., 2016). The online program iTOL (Letunic and Bork, 2016) was used to color tree leaves, allowing easy visualization of which chromosomes each gene was found on. Wheat and *Aegilops tauschii* R gene locations from Ensembl Genomes were used to construct a genomic map using the program Circa (http://omgenomics.com/circa). Clustered wheat R genes were also compared using the program Circoletto (Darzentas, 2010), creating images that display high similarities between genes. Synteny between barley and wheat was determined by aligning chromosome sequences in the program SyMAP (Soderlund *et al*., 2011).

## RESULTS

Approximately half of wheat’s NB-ARC-containing (NBAC) proteins also contained a CC domain, and approximately 75% possessed LRRs **(Table 1)**. Several NBAC proteins contained AP2/ERF, B3, WRKY, and Zinc Finger transcription factor domains. The NBAC proteins also contained kinase, lectin, thioredoxin, and GRAS domains. Of the 2151 NB-ARC-encoding genes, 1505 had NB-ARCs with P-loop, Kinase-2, and GLPL motifs. Among the 1552 proteins that had Leucine-Rich Repeats (LRRs; approximately 72%), 802 contained Coiled-Coil (CC) domains (CNL) and three had Resistance to Powdery mildew 8 (RPW8) domains (RNL). The identified NB-ARC encoding gene accessions are available at the Figshare site (https://figshare.com/s/78d566a9c4cf4902d283). Five of the NB-ARC-encoding genes encoded a Toll/Interleukin-1 Receptor (TIR), with no LRR (TN proteins). NBAC genes formed 547 gene clusters. Clustered genes from each chromosome were compared, resulting in similarity diagrams (Andersen, 2019), illustrating high similarity among clustered genes. Many of the clustered genes possessed greater than 75% similarity within each cluster. While chromosomes from each of the subgenomes were very similar (e.g. chromosomes 1A, 1B, and 1D), differences between the number of clustered genes could have been easily spotted. This was especially noticeable for chromosomes 4A, 4B, and 4D, where 4A contained 57 clusters (involving 119 genes) and 4B and 4D contained four clusters (of eight genes) and three clusters (of seven genes), respectively. It is unknown whether this diversification of NBAC genes in 4A happened prior to the first or second hybridization event in wheat. Since *Triticum urartu* gene locations were not available, a clear understanding of the difference between *T. urartu* chromosome 4 and wheat chromosome 4A could not be reached. **Figure 1** shows a neighbor-joining tree composed of NBAC sequences, specifically those with P-loop, Kinase-2, and GLPL motifs. Nesting patterns in the neighbor-joining tree highlighted additional relationships between clustered genes, displaying which groups arose through tandem duplication, where clustered genes nested together, or segmental duplication, where clustered genes nested separately. Many pairs and triplets of genes from the same cluster nest together on the tree. Due to the close relationship between barley and wheat, synteny between wheat and barley chromosomes was also assessed. Chromosomes of wheat progenitors, such as *Aegilops tauschii*, align with wheat, 1D of wheat matching 1D of *A. tauschii*, and so on. As is visible from the syntenic map (**Figure 2**), the same phenomenon takes place between barley chromosomes (1H-7H) and the three subgenomes of wheat.

**Table 1.**
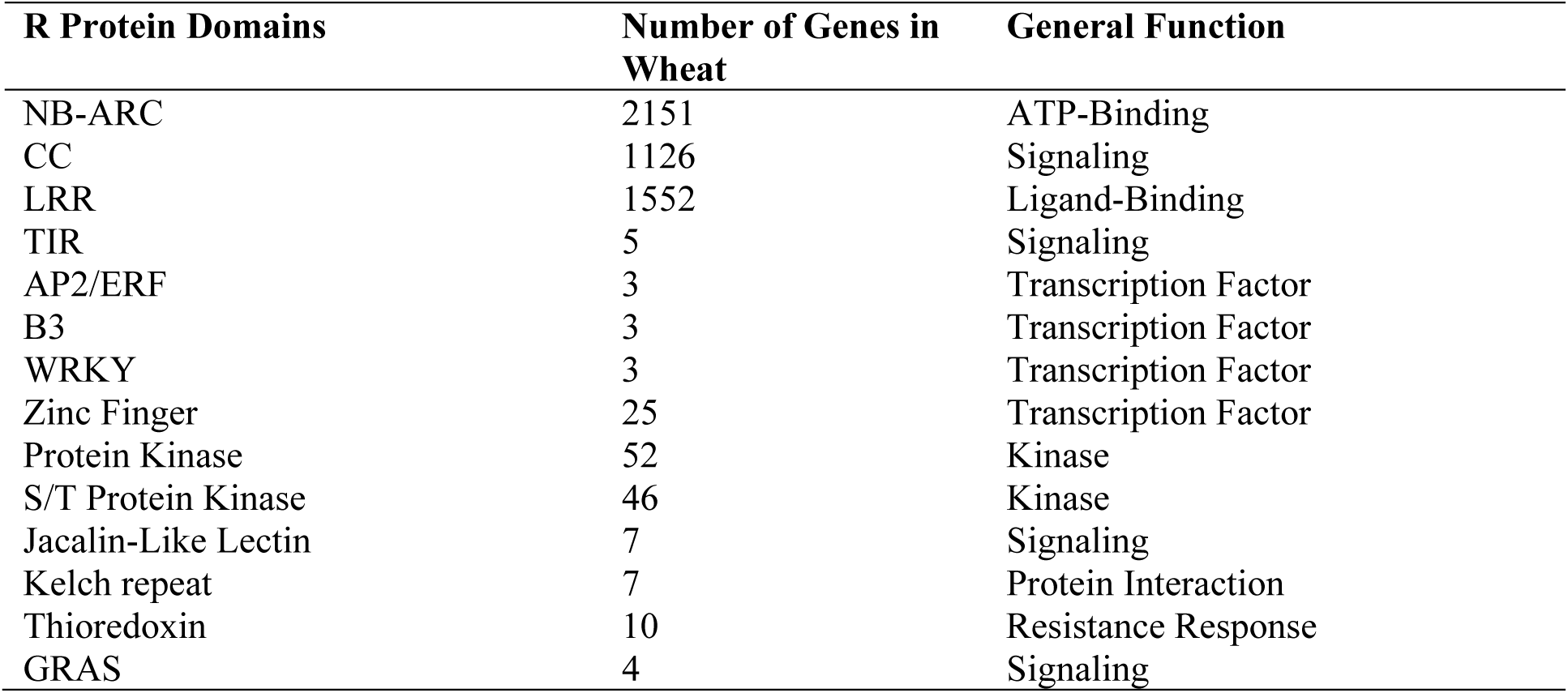
Classification of the identified NB-ARC-encoding genes in wheat.

| **R Protein Domains** | **Number of Genes in Wheat** | **General Function** |
| --- | --- | --- |
| NB-ARC | 2151 | ATP-Binding |
| CC | 1126 | Signaling |
| LRR | 1552 | Ligand-Binding |
| TIR | 5 | Signaling |
| AP2/ERF | 3 | Transcription Factor |
| B3 | 3 | Transcription Factor |
| WRKY | 3 | Transcription Factor |
| Zinc Finger | 25 | Transcription Factor |
| Protein Kinase | 52 | Kinase |
| S/T Protein Kinase | 46 | Kinase |
| Jacalin-Like Lectin | 7 | Signaling |
| Kelch repeat | 7 | Protein Interaction |
| Thioredoxin | 10 | Resistance Response |
| GRAS | 4 | Signaling |

**Figure 1.**
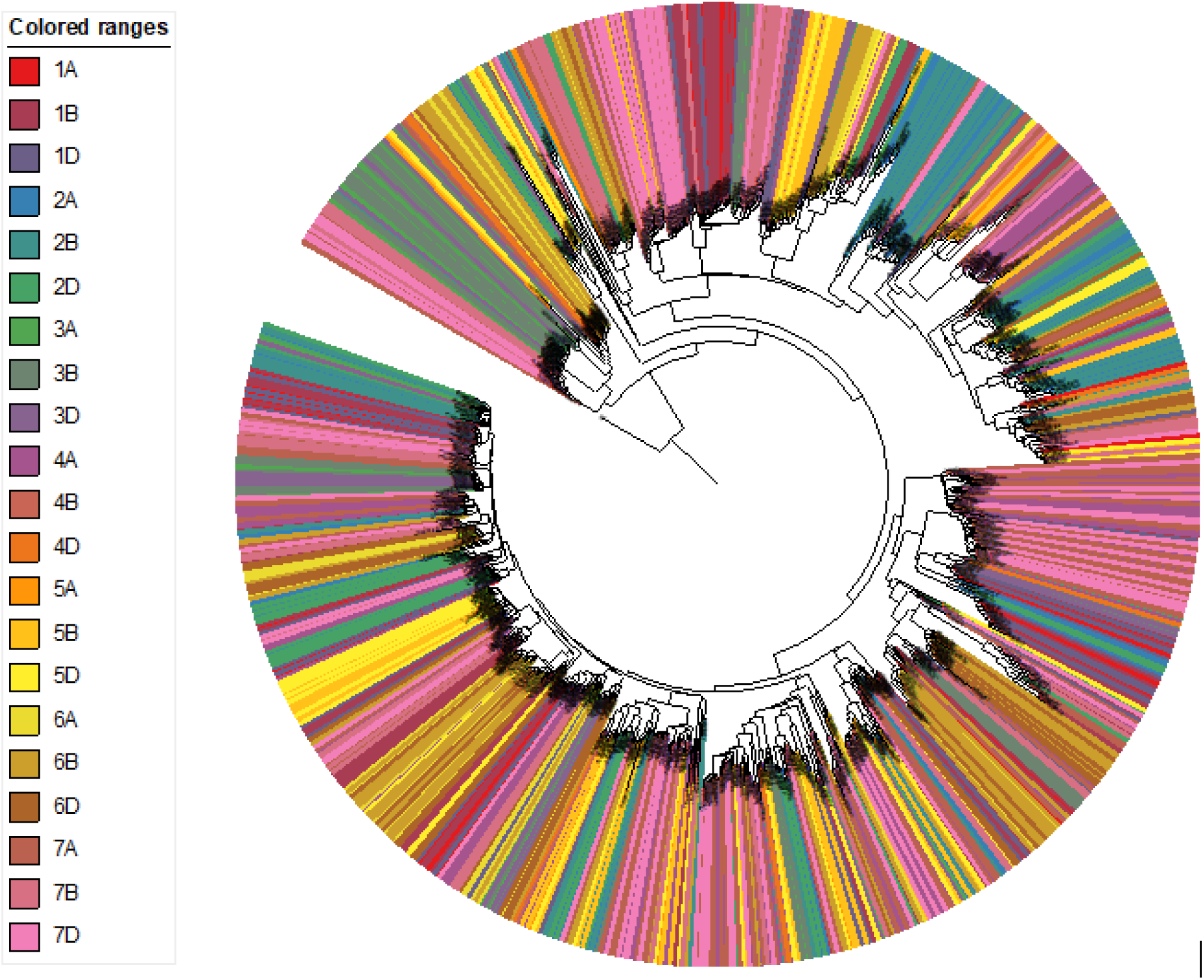
Phylogenetic analysis of clustered NBAC genes. Sequences were included if they contained P-loop, Kinase-2, and GLPL motifs as identified by MEME and if they were found in clusters. This neighbor-joining tree was generated using MEGA 7. Color profiles indicate the chromosomal locations of the gene clusters and A, B and D refer to the source genomes that contributed to the current wheat genome.

**Figure 2.**
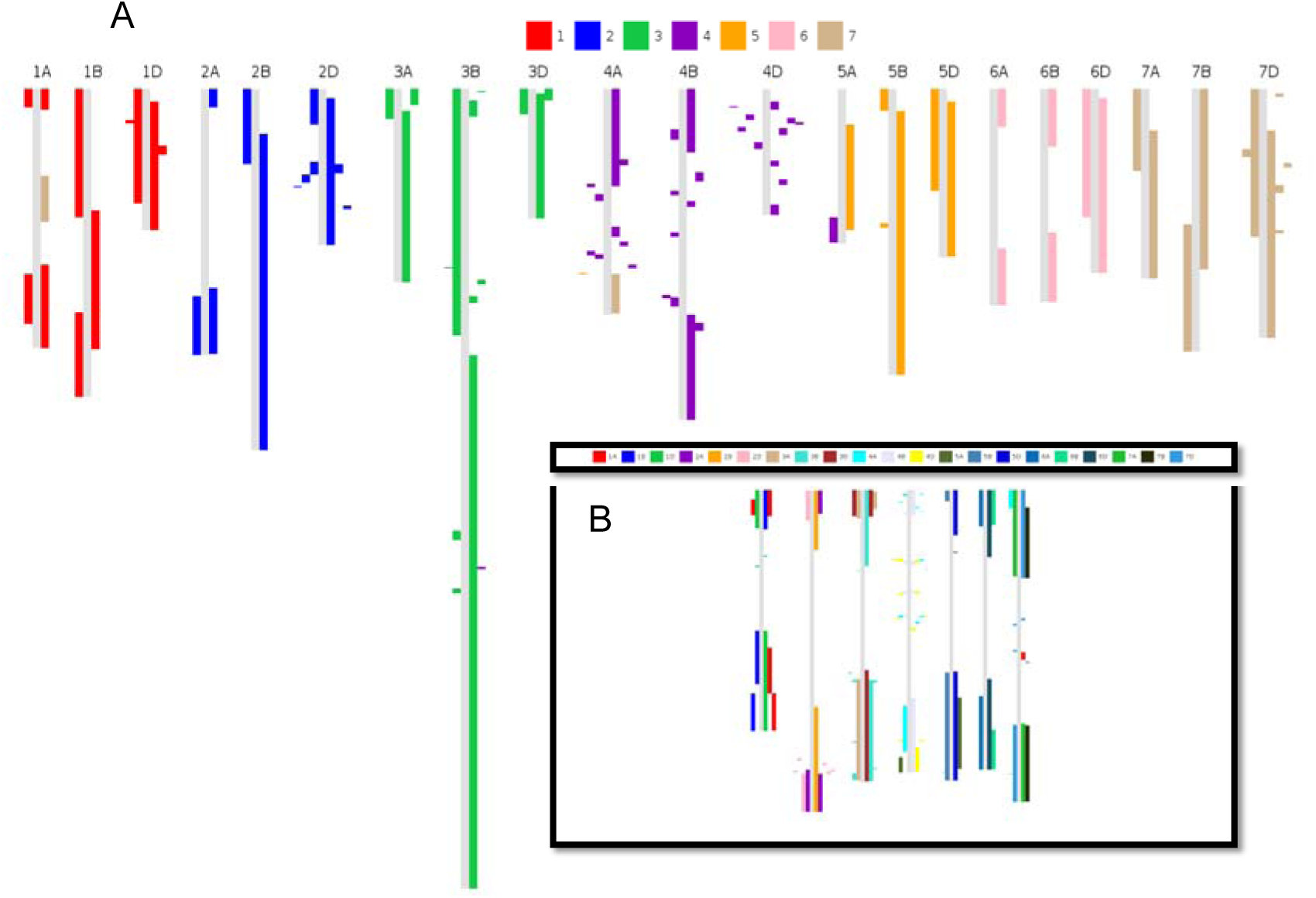
Synteny between chromosomes of wheat and barley. Barley chromosomes were aligned with wheat using the program SyMAP. Barley chromosomes are mapped onto the 21 chromosomes of wheat (A) and the inverse image of wheat chromosomes mapped on the seven chromosomes of barley (B).

## DISCUSSION

This investigation focused on all genes with NB-ARCs because NLR systems have been found that use groups of sensor and helper NLRs to detect and initiate defense responses when pathogenic effectors are present (Jones *et al*., 2016). Not all functional NLRs have all characteristic domains, such as Pb1 (Hayashi *et al*., 2010). Excluding genes that do not contain particular domains or motifs may not include important genes that assist with resistance responses. Therefore, proteins lacking CC or LRR domains may also contribute to resistance responses, especially those with additional domains that are involved in signaling (Baggs *et al*., 2017). The distribution of NBAC genes across wheat chromosomes concurs with previous studies in barley and foxtail millet, where R genes were also found in clusters in extra-pericentromeric regions of chromosomes (Andersen *et al*., 2016; Andersen and Nepal, 2017). Unequal crossing over between chromosomes as a mechanism for duplication likely explains the formation of these clusters, which then allows for their diversification. Previous studies have highlighted this explanation for the location of the quickly evolving genes (Marone *et al*., 2013). *A. tauschii* and *H. vulgare* share a similar pattern as wheat (Andersen, 2019), with a similar number of R genes located at the ends of chromosomes. Analyses of the clusters of NLR genes revealed many genes with high sequence similarity (Andersen, 2019), indicating their origin by tandem duplications, especially those that are only a few hundred nucleotides away from each other and share >90% similarity. Through tandem duplication, wheat NLR genes may have diversified to respond to rapidly evolving and perhaps closely related pathogens. Many pathogens possess diverse races, such as the pathogen *Pyrenophora tritici-repentis* with eight races (Abdullah, 2017), and race-specific (vertical) resistance has been identified to wheat pathogens, such as powdery mildew (Bourras *et al*., 2015). This type of resistance involves one or two genes, as opposed to horizontal resistance, which involves multiple genes (quantitative) and provides resistance to many pathogens. Horizontal resistance may include other signaling factors and types of receptors, relying only partially on NLRs (Kushalappa *et al*., 2016).

While it can be inferred that similar genes that are close to each other on a chromosome are likely due to tandem duplications, several genes were dissimilar and close together. This can be seen in **Figure 1**, where, while many clustered genes nest together, providing visual evidence of tandem duplications with closely clustered genes nesting together, many do not nest together. This phenomenon has two major explanations: 1) the tandem duplication took place long ago in evolutionary history and has had time to diversify greatly, or 2) a segmental duplication took place, causing the gene to become located next to another R gene or R gene cluster. R genes are highly diversified in plants, many plants possessing hundreds of them, indicating that these R genes originate from very ancient precursors. Ancient tandem duplications would have much time to diversify, especially if particular selective pressures are put upon the ancestors of modern species. However, segmental duplications cannot be discounted due to the presence of genes in some clusters that are highly similar to genes in other clusters (Leister, 2004). In these cases, transposable elements may play some role in movement of these genes around to other chromosomes or distant locations on the same chromosome (Kim *et al*., 2017).

Barley and the progenitors of wheat diverged only approximately 8-9 million years ago (Middleton *et al*., 2014). Therefore, barley provides an excellent partner for wheat synergistic comparison. Both barley and wheat experienced artificial selection since both have been grown for food production since the agricultural revolution approximately 10,000 years ago. Wheat differs from barley in that it is an allohexaploid resulting from hybridization of three species, each containing seven pairs of chromosomes to total 21 pairs, while barley remained diploid with only seven pairs of chromosomes. The wheat genome, consisting of A, B, and D subgenomes, maps to the barley genome (H), with wheat chromosomes 1A, 1B, and 1D containing much synteny to 1H of barley. The syntenic map (**Figure 2**) displays this similarity between genomes. NLR gene clusters in these syntenic blocks were investigated to see which duplications took place before the wheat-barley divergence. Instances exist where barley possesses duplicated genes that remained individuals in wheat, and vice versa. The similarity between wheat and *A. tauschii* is much closer, since *A. tauschii* contributed wheat’s D subgenome only a few thousand years ago. While the A subgenome progenitor, *Triticum urartu*, has limited genomic availability, future studies may be able to assess differences between the NLR gene architecture in the two genomes. *A. tauschii* and barley provide excellent comparisons with wheat due to the relatively short period of time since their divergence. The similarities between R genes in wheat relatives show that the highly diverse family of R genes is necessary for survival, whereas the differences in number and phylogeny point to differences in selection pressure that these species each face.

## CONCLUSION

In this study, clustering of R genes in wheat has been described, as compared to its progenitors and barley, a close relative. Gene similarities within clusters were assessed, showing that tandem duplication explains some of the diversification among R genes, along with segmental duplication and possible action by transposable elements. Wheat possesses 2151 NB-ARC-encoding genes, with many of those encoding the domains associated with functional NLRs that likely function as receptors, detecting pathogenic effectors. In wheat’s 21 chromosomes, 547 clusters were found, with many of them containing highly similar genes. Future research should seek to functionally classify which pathogens these proteins trigger responses to and if duplications can be associated with known events in wheat’s evolutionary history. Additional genomic data on *Aegilops speltoides*, a relative to the contributor of wheat’s B subgenome, as well as availability of data for *Triticum urartu*, contributor of wheat’s A subgenome, will allow for more thorough analysis of the evolution of disease resistance genes in wheat.

## ACKNOWLEDGEMENTS

Support for this research came from the USDA-NIFA hatch Projects to M. Nepal (SD00H469-13 and SD00H659-18). The paper represents Lauren Lindsey’s undergraduate capstone project integrated with Ethan Andersen’s PhD dissertation research at South Dakota State University.

